# Eccentric contraction response of stimulated skeletal muscle fascicle at the various strain rates and stimulation timing

**DOI:** 10.1101/2023.04.26.538494

**Authors:** Dat Trong Tran, Liren Tsai

**Affiliations:** Department of Mechanical Engineering, National Kaohsiung University of Science and Technology, Kaohsiung, Taiwan; School of Transportation Engineering, Hanoi University of Science and Technology, Hanoi, Vietnam

**Keywords:** Eccentric contraction, strain rate, muscle stimulation, muscle injury.

## Abstract

Muscle injuries are the most common sports injuries, and it is often observed in eccentric contraction. There are many factors that could influence the severity of muscle injuries, including strain and strain rate. This study evaluated the interaction of these two factors on the biomechanical properties of the muscle-tendon bundle and their role in injuries. A Hopkinson bar system, an MTS machine and an electrical pulse generator were utilized to collect eccentric contraction response data of over 150 frog muscle-tendon samples at strain rates ranging from 0.01 to 300 s^-1^. The results have shown that the maximum stress has increased and peaked at about 150 s^-1^. That peak value has then maintained at the following strain rates. While Young’s modulus of stimulated samples reduced as the strain rate changed from 50 to 300 s^-1^. That trend was in contrast to unstimulated muscle bundles. In addition, strain rate has significantly influenced tendon-muscle bundle fracture. Samples tend to rupture at a minor strain of about 3.5 % with strain rates over 200 s^-1^. Because of the increasing stiffness of the muscle area at high strain rates, increased strain in the tendon region resulted in frequent injuries in the tendon area. On the other hand, a maximum-stress reduction was detected when the muscle bundles were stimulated at muscle strain greater than 0.2. The results showed that improper timing of stimulation could increase muscle injury.

## 1. Introduction

Muscle injuries often occur during sports activities [1–3], and researchers have conducted numerous studies to determine their causes by examining the biomechanics of muscles during movement. Those studies have indicated muscle injury occurs in response to forcibly passive stretching, and passive muscle tension is strongly related to function. However, the injuries often occur during eccentric contraction [4–7]. An eccentric muscle contraction is observed when a tension force applied to the muscle exceeds the momentary force generated by muscle stimulation, resulting in forced lengthening of the muscle-tendon system while contracting [8]. Eccentric contraction often occurs in downhill running, especially during intense activities [9]. Gabbe et al. reported that most muscle injuries in community-level Australian football occurred during running or sprinting (about 80%), while only 19% occurred in kicking [10]. Force does not seem to be the determining factor in the injury. This point of view was also stated in Hui Liu’s report [11]. Therefore, correctly understanding the biomechanics of muscles is essential for predicting and preventing injuries during vigorous exercise.

The biomechanical behavior of muscle tissue materials, including active, passive contraction and eccentric contraction, has been invested over the years at all scales, from fiber, bundle, and fascicle scales to whole muscle-tendon architecture. Passive biomechanics depends on many factors, such as temperature, strain, strain rate, and scale of muscle bundle, where strain rate plays an important role [12–18]. Most studies indicate that tendon muscles’ stress and elastic modulus increase at higher strain rates. In active contraction and eccentric biomechanism, the strain rate or contraction velocity is always an important factor affecting the properties of the muscle bundle [12–21]. Therefore, they are parameters that should be considered when studying the load capacity and risk of the tendon–muscle bundle injury. However, many studies have shown that muscle strain significantly affects injury rates [11, 16, 22]. The greater muscular elongation velocities increase the severity of muscular strain injury in the extensor digitorum longus muscles, but only during large muscular strains[23–26]. Therefore, it is difficult to separately evaluate each factor, such as strain and strain rate, in studying the properties and mechanism of injury in the tendon-muscle bundle.

In this study, we invested the passive and eccentric contraction responses of the whole muscle-tendon-bone structure with the experiments of over 150 frog muscle-tendon samples at different strain rates from 0.01 to 300 s^-1^. On the other hand, the muscle bundle will be stimulated at various strains as they are tested at low strain rates from 0.01 to 0.5 s^-1^. In contrast, the muscle bundle will be applied to the stimulation to generate a large isometric contraction at two distinctive lengths (0% and 10% initial length) before an eccentric contraction is applied at the strain rate from 50 to 300 s^-1^. The results will be used to evaluate the influence of deformation and strain rate on the eccentric contraction property. Simultaneously, the impact of factors including stimulus, strain and strain rate was assessed on the risk of muscle injury during sports activities.

## 2. Material and method

The frogs of similar size were acquired at a local market and immediately transferred to the laboratory to dissect the frog’s semi-tendinosis muscle-tendon bundles. The samples were preserved in 0.9% saline solution after dissecting and during the property test. In addition, the samples were moistened with tissue impregnated with saline solution after being installed on the clampers. A total of over 150 specimens of 40 ± 3 mm in length and roughly 4 ± 0.2 mm in diameter were prepared for this experiment.

Active muscle contraction is affected by many parameters of the electrical stimulation pulse, such as voltage, frequency and duty. However, it could be confirmed that muscle contraction increases with the amount of released calcium (higher frequency will release more calcium); however, to a saturation stage, more Ca2+ release will not result in increased force [20, 27, 28]. To get a large muscle contraction, we experimented with selecting the parameter of the stimulation pulse. After some tests with GW INSTEK AGF-3051 and PPH-1503 generators, a pulse with a frequency 40Hz, 50% duty and amplitude of 10V was selected for this study (about 30 mA and inter-pulse interval of 12.5 ms). This voltage and frequency ensured that recruited responsive motor units were as much as possible and the muscle fatigue was small [20, 29–32]. Needle electrodes were used to deliver pulses to the muscle bundle, and the distance from the bond to the electrodes was about 8 mm [33].

LLOYD Friction Testing machine made by AMETEK (Figure 1) was utilized for experiments at low strain rates from 0.01 to 0.5 s^-1^. On the other hand, two clamps (Figure 1) were made by 3D printing to secure the samples at both ends and the center of the device shaft without sliding. In addition, the clamps were fabricated with PLA material possessing a large elastic modulus and cross-section, which ensured minimal deformation during the test and prevented any effect on the accuracy of the results.

**Fig. 1.**
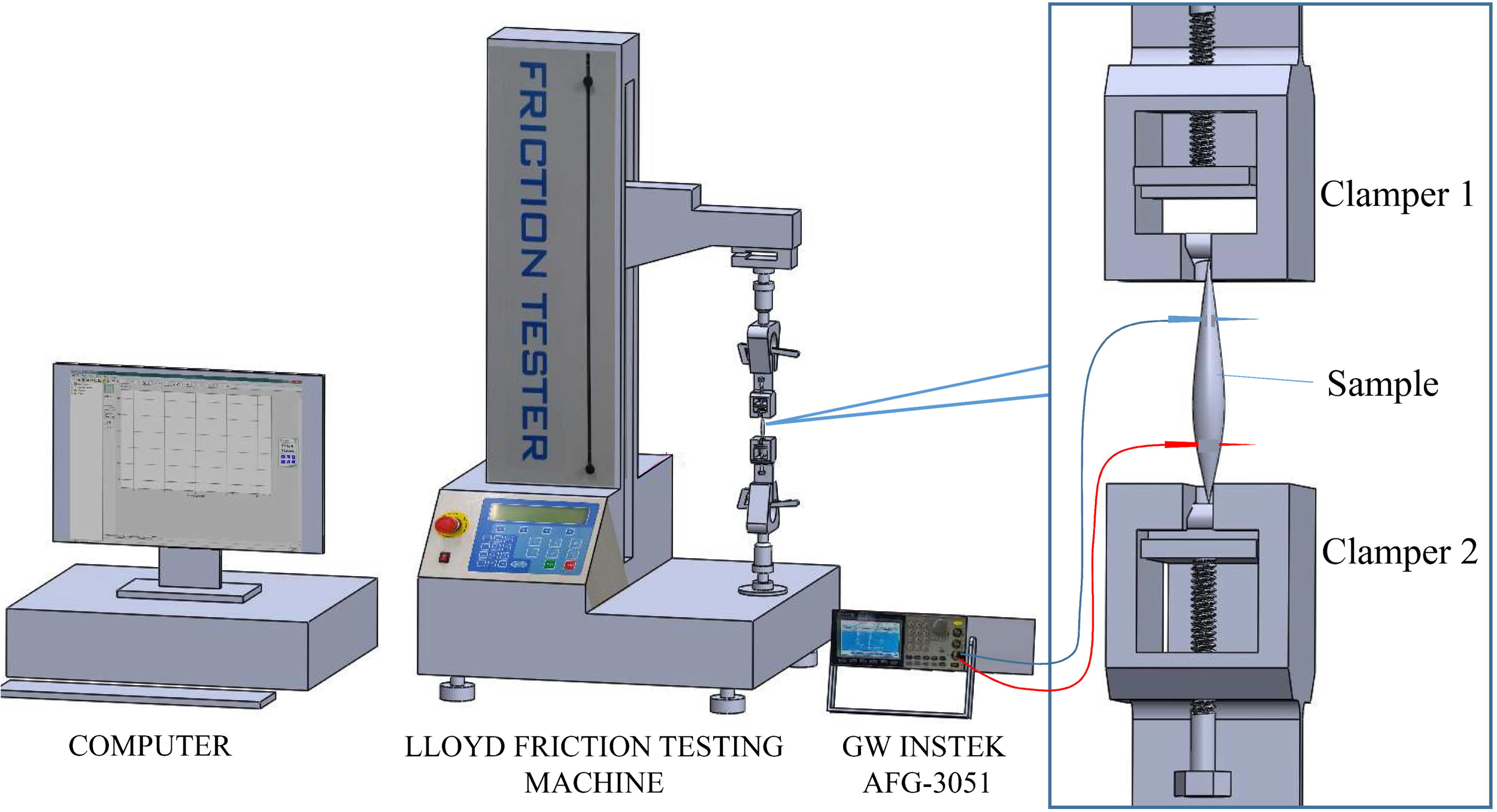
Structure of the experiment at low strain rates.

To begin the experiment, the samples were mounted onto clamps and pulled to 0.2 N to establish their initial length. The electrodes were then installed with the experiments using the stimulation. Each strain rate value was tested on 8 to 9 samples, all of which were elongated from their original length to 100% deformation. Firstly, the unstimulated muscle-tendon bundles were subjected to property testing at strain rates from 0.01 to 0.5 s^-1^. For tests involving stimulated muscle bundles, stimulation was applied at strain rates of 0.02 to 0.03 and maintained until the samples were fully broken at 100% deformation in the 0.01 and 0.1 s^-1^ strain ranges. However, the test process happened in a short-time range at 0.5 s^-1^ strain rate; the stimulation was generated until the maximum active muscle force was reached, which took approximately 2 seconds. Afterwards, the tension process would be applied. The muscle bundle’s mechanical properties were assumed to remain unaffected as the difference in muscle strain when the stimulus was applied was negligible (Table 1).

**Table 1.** Experimental conditions were determined based on the muscle bundle’s strain when the stimulus was generated.

| Initial strain applied stimulation at the low strain rate<br>(From 0.01 to 0.5 s<sup>-1</sup>) |  |  | High strain rate<br>(From 50 to 300 s<sup>-1</sup>) |  |
| --- | --- | --- | --- | --- |
| Strain rate<br>0.01 s<sup>-1</sup> | Strain rate<br>0.1 s<sup>-1</sup> | Strain rate<br>0.5 s<sup>-1</sup> | L0 group<br>(0% strain) | L10 group<br>(10% strain) |
| 2 % to 3 % | 2 % to 3 % | 0% | 0 % | - |
| 20, 30 and 40% | - | - | - | 10% |

In addition to the aforementioned tests, we conducted experiments at a strain rate of 0.01 s-1 to analyze the effect of stimulated timing on muscle function. The muscle was stimulated at 20%, 30%, and 40% strains until reaching 100% strain. We chose a strain rate of 0.01 s-1 for this test as it provided sufficient time to apply the required stimulation precisely. These tests aimed to assess the impact of nerve stimulation timing during an eccentric contraction on the muscle. To ensure the accuracy of the results, each sample was tested only once, preventing the previous testing process from affecting the subsequent results.

In experiments at high strain rates from 50 to 300 s^-1^, the mechanical properties of stimulated skeletal muscle fascicles were collected by the Hopkinson bar system (Figure 2), which combined three 6061 aluminium bars: a 400 mm striker bar, a 2000 mm incident bar, and a 1000 mm hollow transmitted bar. Four micro sensors have been arranged on the incident and transmission bar to measure the deformation of the bars. A collision between Stricker and the Incident bar was produced by the striker’s movement driven by a pneumatic cylinder. The deformation wave in the incident bar (ɛ_i_) has propagated to the other end of the bar where the sample was clamped. A part of this pulse has passed through the sample to the transmission bar (ɛ_t_), and another has passed back to the incident bar (ɛ_r_). Interactions between impulses produce deformation in muscle-tendon bundles. Strains on the bars were calculated based on signals from the sensors collected by an oscilloscope. The mechanical properties, including strain, stress and strain rate 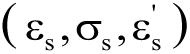, are calculated from the following equations based on one-dimensional stress wave theory [34–37]:

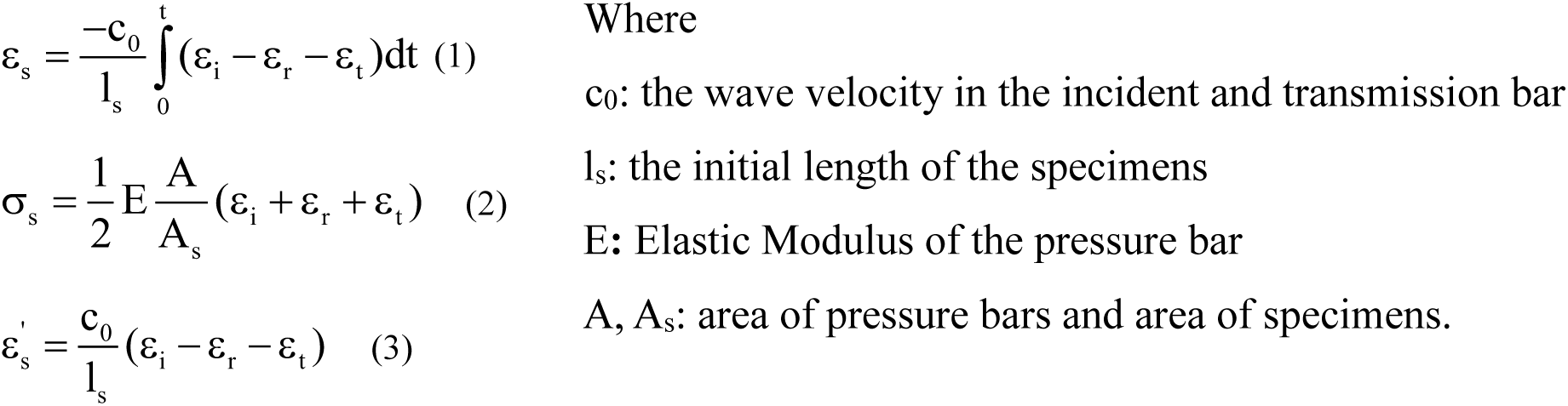

**Fig. 2.**
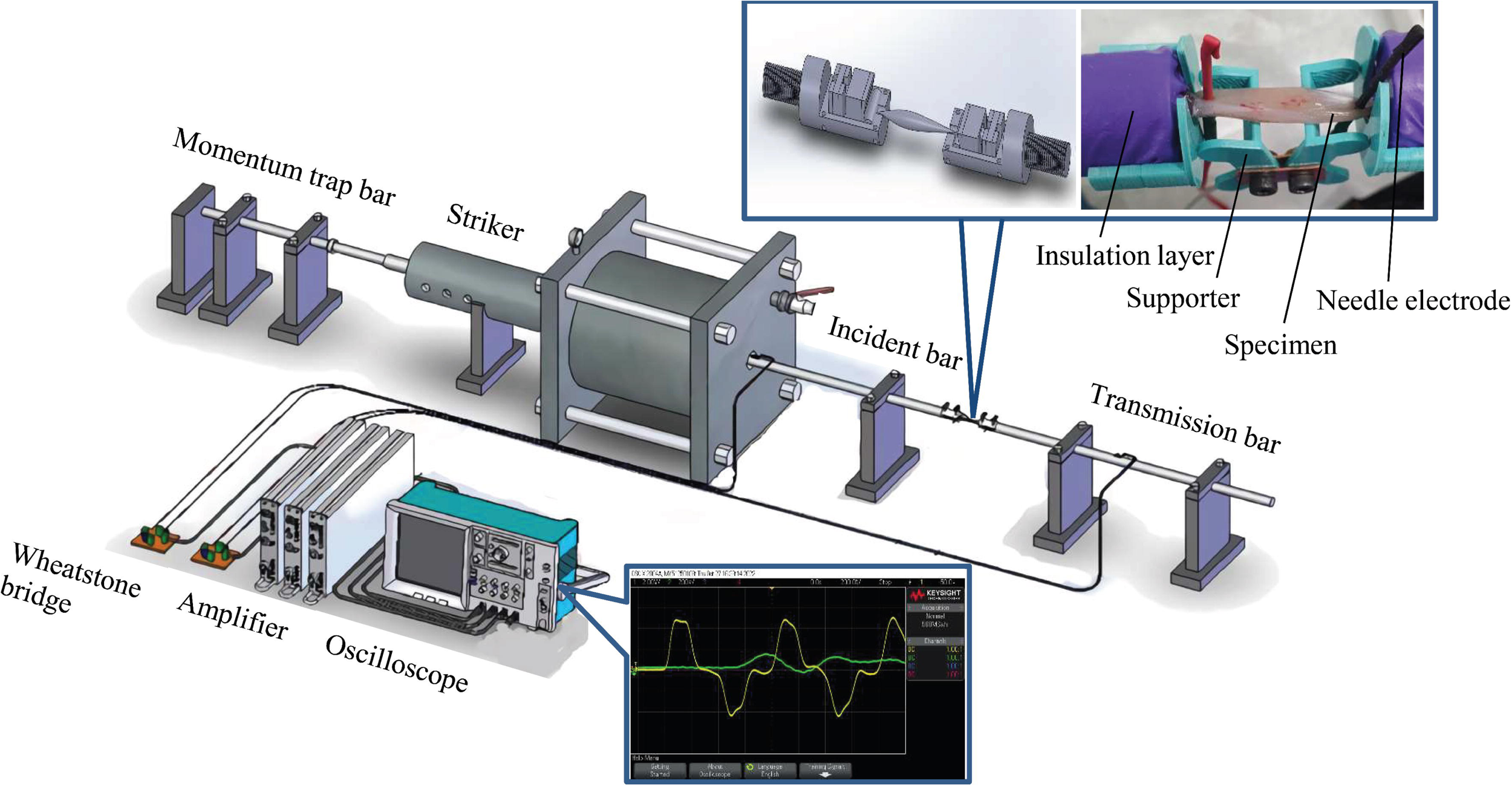
Structure of the experiment at the high strain rate.

This study divided samples into two groups at high strain rates. In group 1 (L0), the samples were clamped on the rod ends, and the needle electrode was mounted on samples about 8mm from the bone (Figure 2). A supporter was used to maintain the muscle bundles at their original length. When the muscle bundles were stimulated, an isometric contraction occurred before the Hopkinson bars system was activated to create an eccentric contraction in the muscle bundles, which occurred about 3 seconds later. The effect of strain applied a stimulation could be accessed at the low strain rate range. However, some stimulation conditions could show different results at the high strain rate range. So in group 2 (L10), samples were applied an isometric contraction at length greater than 10% before being pulled to produce an eccentric contraction. Both groups tested the properties of skeletal muscle bundles at five strain rates, from 50 to 300 s^-1^. For each strain rate, eight samples were tested. In addition, specimens were used only once to ensure optimal muscle contraction. This procedure was performed the same in both groups.

## 3. Results

Figure 3 illustrates the response of eight unstimulated muscle-tendon specimens at a strain rate of 0.01 s^-1^. The data was used to create an average curve using the moving average algorithm, represented by the solid black line. Similar property tests were conducted for the specimens at strain rates of 0.1 and 0.5 s^-1^, and three average stress-strain curves are shown in Figure 4. These curves are similar to previous studies, with a slack area observed at a strain of 0.2. After this region, the stress increases rapidly and peaks at a strain of 0.50 to 0.62, consistent with previous findings [38–42]. The sensitivity of the muscle-tendon bundle to strain rate is evident as Young’s modulus and maximum stress increase with increased strain rate.

**Fig. 3.**
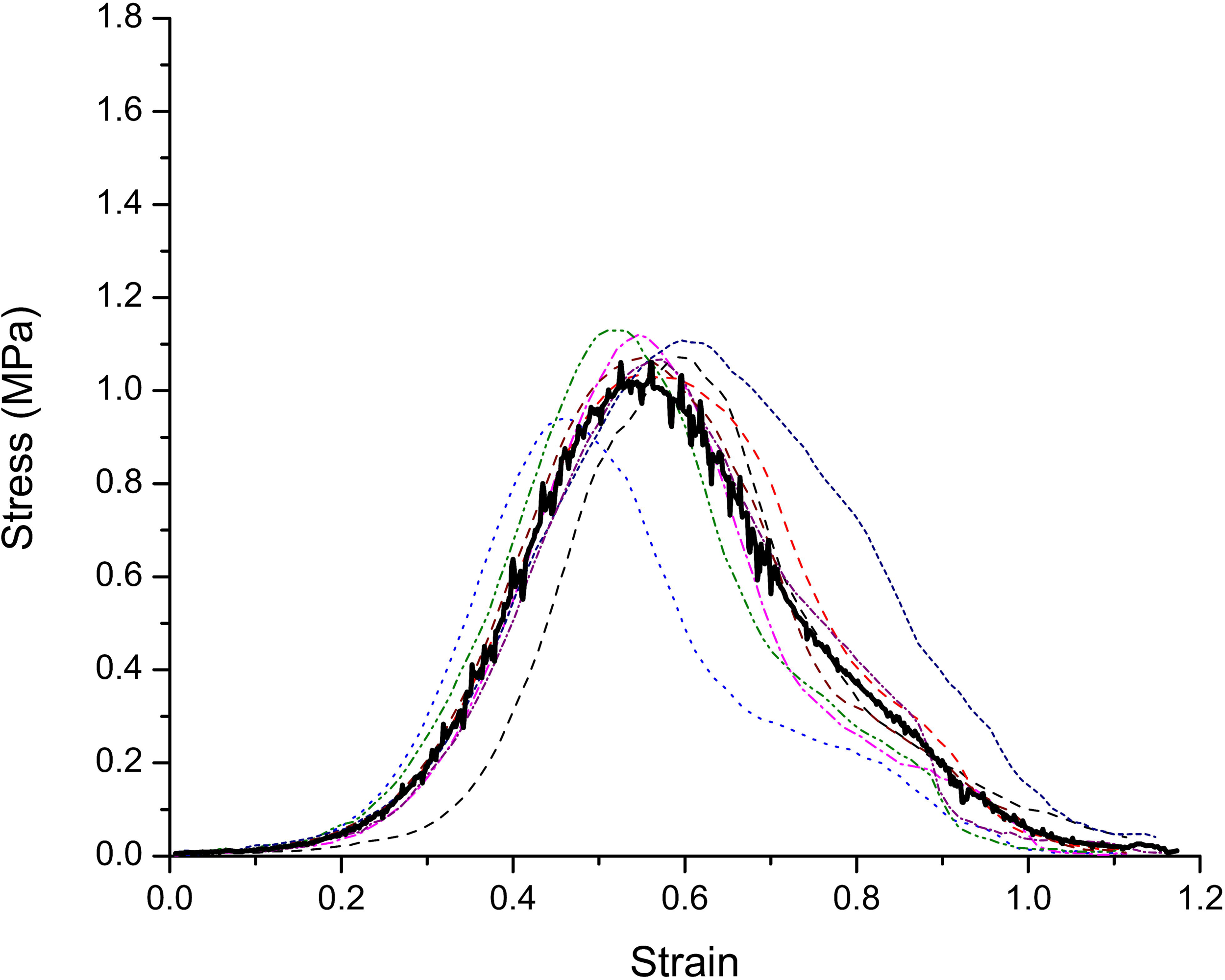
The stress-strain curves of skeletal muscle at strain rate 0.01 s^-1^ without a stimulation

**Fig. 4.**
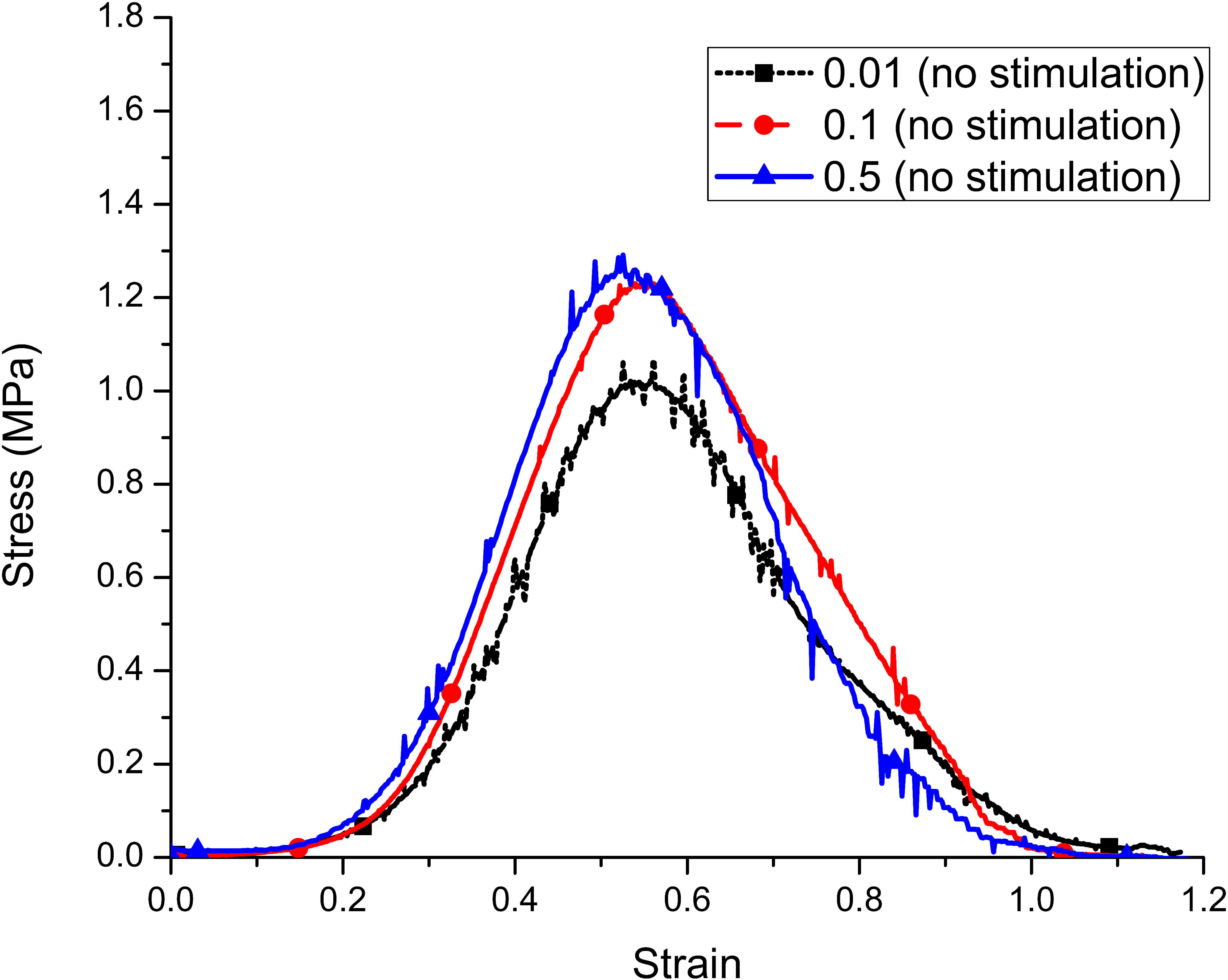
The average stress-strain curves of unstimulated skeletal muscle at three various strain rates

The study investigated the eccentric contraction response of skeletal muscle at 3 similar strain rates with the stimulations outlined in the method section. At strain rates of 0.01 and 0.1 s^-1^, the stimulation pulses were immediately applied after the samples were tensioned. For the 0.5 s^-1^ strain rate, stimulations were applied just before the samples were pulled to accommodate the shorter test duration. Nine specimens were tested at each strain rate (Fig. 5), and the mean curves were also calculated (solid black line). In stimulated samples, the stress created below 0.05 strain was considered active stress (the passive stress was insignificant in this strain range, as shown in Fig. 4 and Fig. 6). The eccentric contraction stress also increased with the increase of strain and was equal to the sum of the active and passive stress components. These findings were consistent with previous research[42–44]. On the other hand, the research found that the active stress component in an eccentric contraction of skeletal muscle at 0.2 strain significantly increased about twice from 0.4 to 0.8 MPa in stimulated muscle bundles when the strain rate increased from 0.01 s^-1^ to 0.1 s^-1^ (Figure 6). Additionally, a slack region was observed between 0.1 to 0.3 strain before the stress continuously increased to the maximum stress value. That concluded that the muscle contraction velocity and strain rate significantly rose the active contraction component of an eccentric contraction[21, 45–48]. However, the slack region was not observed at the average curve and curves in the experiment at the 0.5 s^-1^ strain rate. The tendon-muscle response showed a linear region from the start to the maximum stress point.

**Fig. 5.**
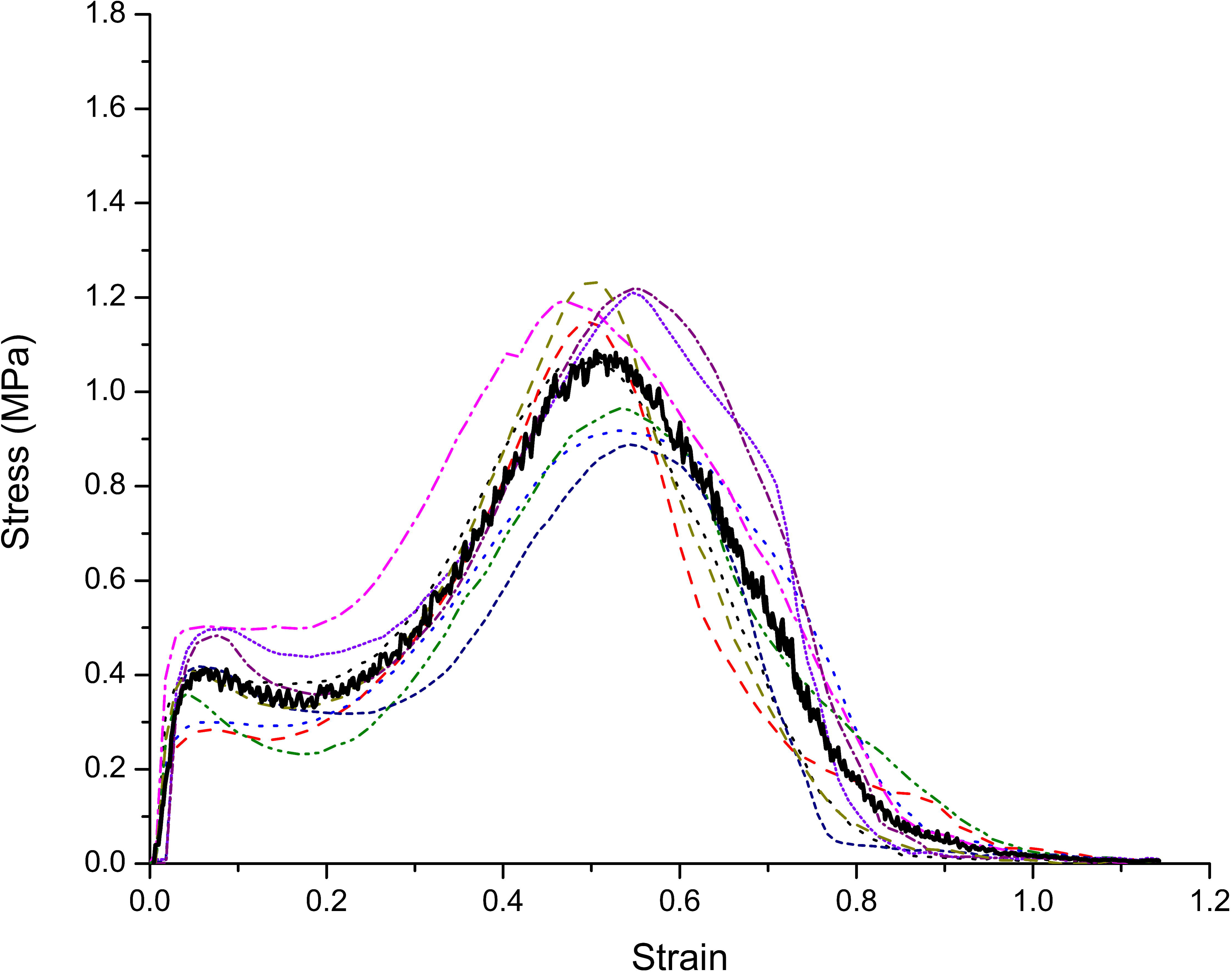
the stress-strain curves of skeletal muscle at strain rate 0.01 s^-1^ with the stimulation applied at the beginning of tension.

**Fig. 6.**
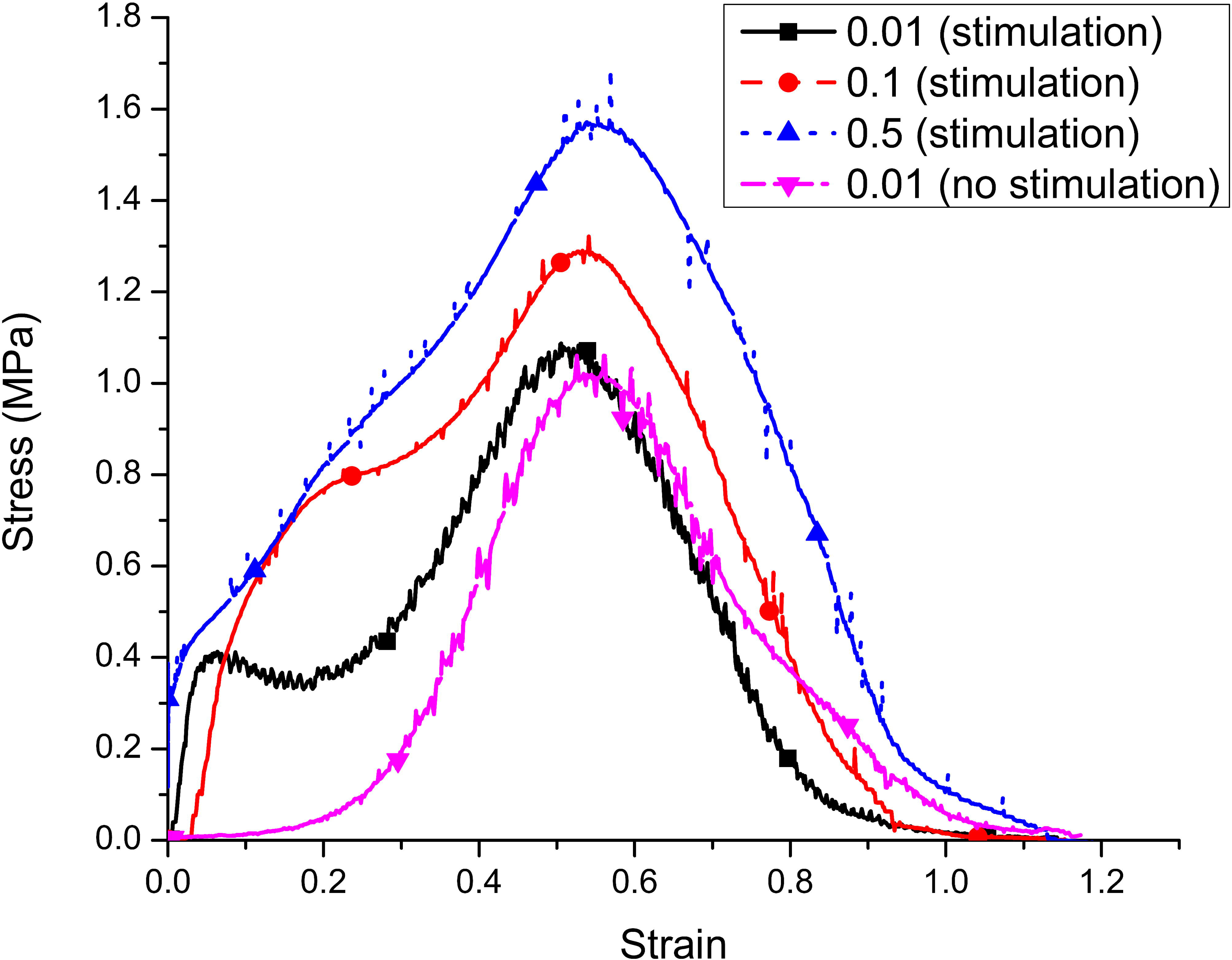
the average stress-strain curves of stimulated skeletal muscle at three various strain rates.

Figure 8 illustrates the average stress-strain curves of stimulated muscle-tendon specimens (L0 group) at the original length in 5 strain rate ranges from 50 to 300 s^-1^. Each curve in Figure 8 was constructed from the property data of 8 samples tested over the same strain rate range as in Figure 7. The slack area was not observed as a property of unstimulated muscle-tendon bundles [38–41]. Muscle bundles demonstrated linear behavior at strain rates above 200 s^-1^. On the other hand, The maximum stress also increased with the increase of strain rate at the scope of the strain rate from 50 to 200 s^-1^, while stress slightly reduced at the strain rate of 300 s^-1^, with the strain changing from 1 to 5.5 percent. This eccentric stress increase was similar to the strain rate’s effect on passive muscle stress, as reported in previous research[43, 44]. In this group, all specimens at strain rates of 200 s^-1^ and 300 s^-1^ were broken entirely in the tendon-muscle connection area or closer to the tendon region [49]. In addition, the samples showed a decrease in Young’s modulus from about 78MPa to 35MPa. That trend was contrasted with the unstimulated samples (Figure 9 and Table 2). However, Young’s modulus of stimulated samples was still higher than the unstimulated samples in both strain rate ranges as compared to previous research [42] (about 19 to 30Mpa).

**Fig. 7.**
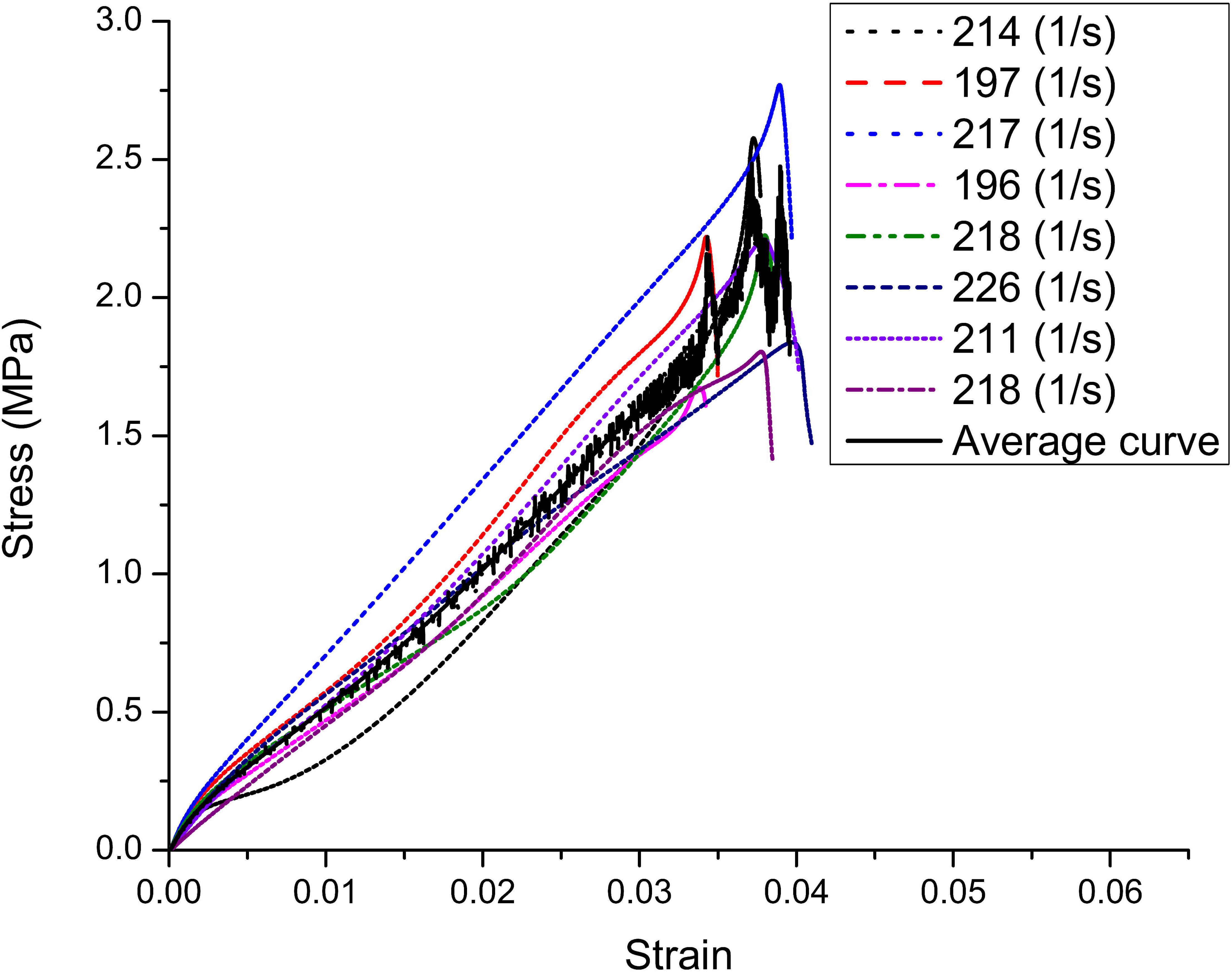
The stress-strain curves of 8 muscle-tendon specimens at the 200 1/s strain rate range stimulated at the original length (L0).

**Fig. 8.**
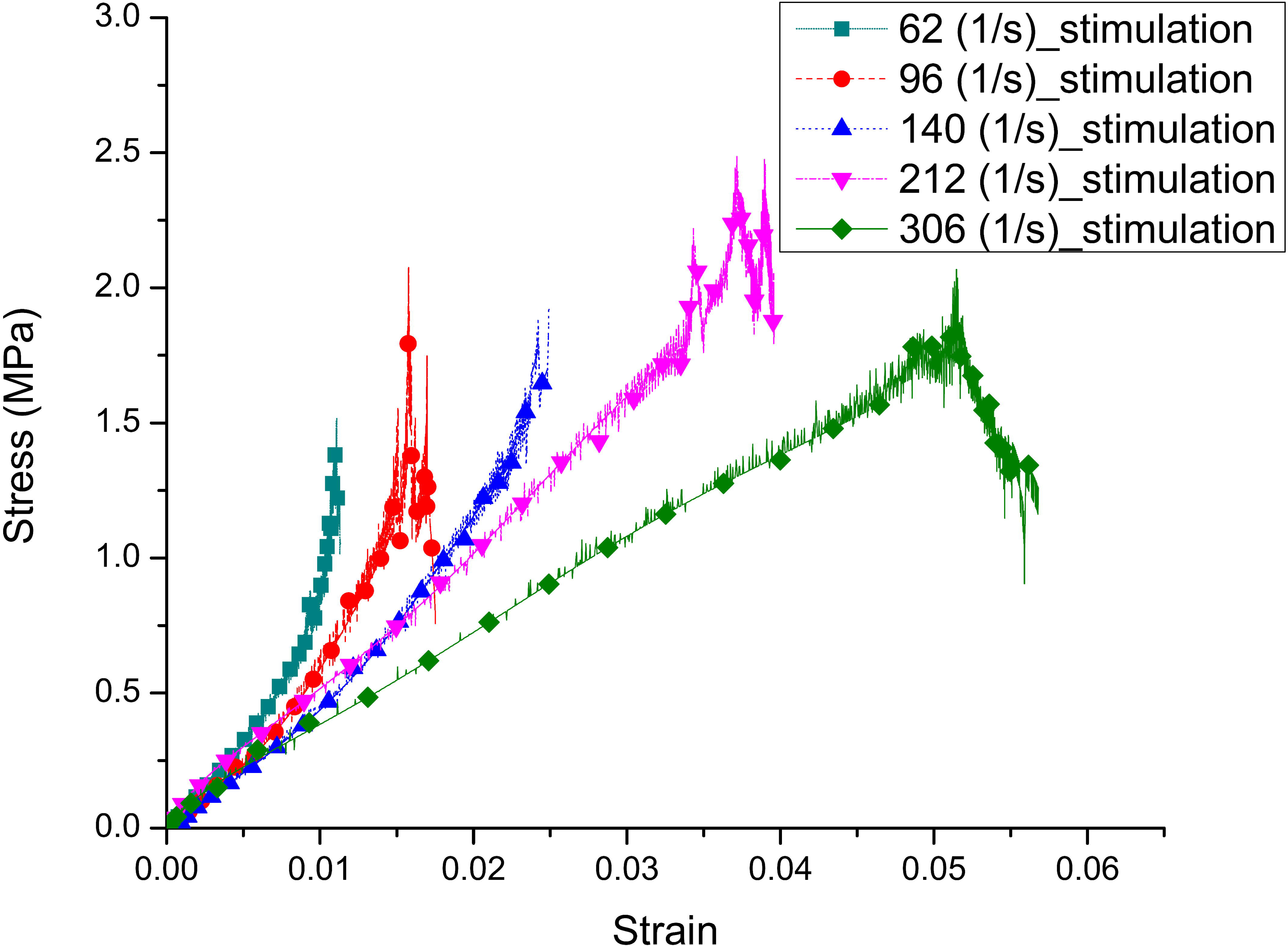
The average stress-strain curves of muscle-tendon specimens stimulated at the original length (L0).

**Fig 9.**
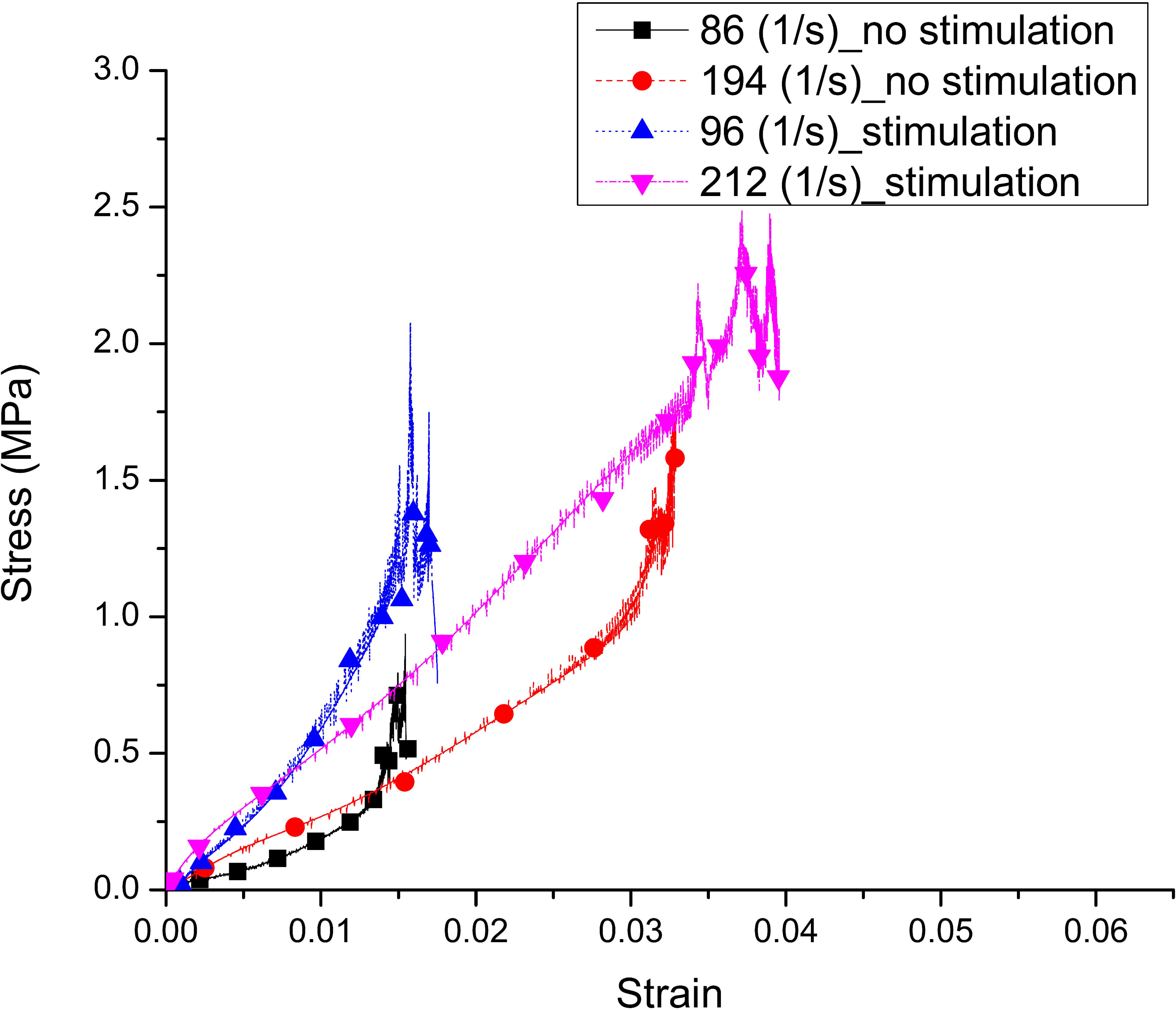
The average stress-strain curves of stimulated and instimulated muscle-tendon specimens stimulated [42].

**Table 2.** Young’s modulus of muscle-tendon bundles at high strain rates

| Strain<br>rate range<br>($\text{s}^{-1}$) | L10 group (stimulation) | L0 group (stimulation) | No stimulation |
| --- | --- | --- | --- |
|  | Young's<br>modulus (MPa) | Young's<br>modulus (MPa) | Young's<br>modulus (MPa) |
| 50 | 66.24 | 78.07 |  |
| 100 | 50.66 | 67.16 | 19.71 |
| 150 | 55.46 | 58.59 |  |
| 200 | 44.04 | 51.85 | 30.39 |
| 300 | 44.37 | 34.60 |  |

The values of the maximum stress and the strain corresponding to maximum stress in the experiments were summarized in Figures 10 and 11. It showed that the maximum stress in unstimulated samples was sensitive to strain rate at low strain rates (the maximum stress increased from 1.06 to 1.34 MPa)[12–15]. The stimulations also dramatically affected the break stress. The maximum stress was increased from 1.09 to 1.62 MPa. The effect of the electric pulse was more evident at the strain rate of 0.5 s^-1^, where the mean maximum eccentric stress increased by about 21% more than the passive stress. The influent of strain rate on maximum eccentric stress was continuously observed at higher strain rates. However, the maximum stress increase was gradually reduced at the higher strain rate (Figure 11), and the highest value was about 2 MPa. That trend was confirmed in the previous research [19–21]. Although affecting the maximum stress, the stimulation pulses did not affect the breaking strain of samples at the low strain rate, and all experiments illustrated that the muscle bundles were broken in the range of 50 to 55%. The effect of stimulation on the strain of samples was also insignificant at the high strain rate, showed in Figure 9.

**Fig. 10.**
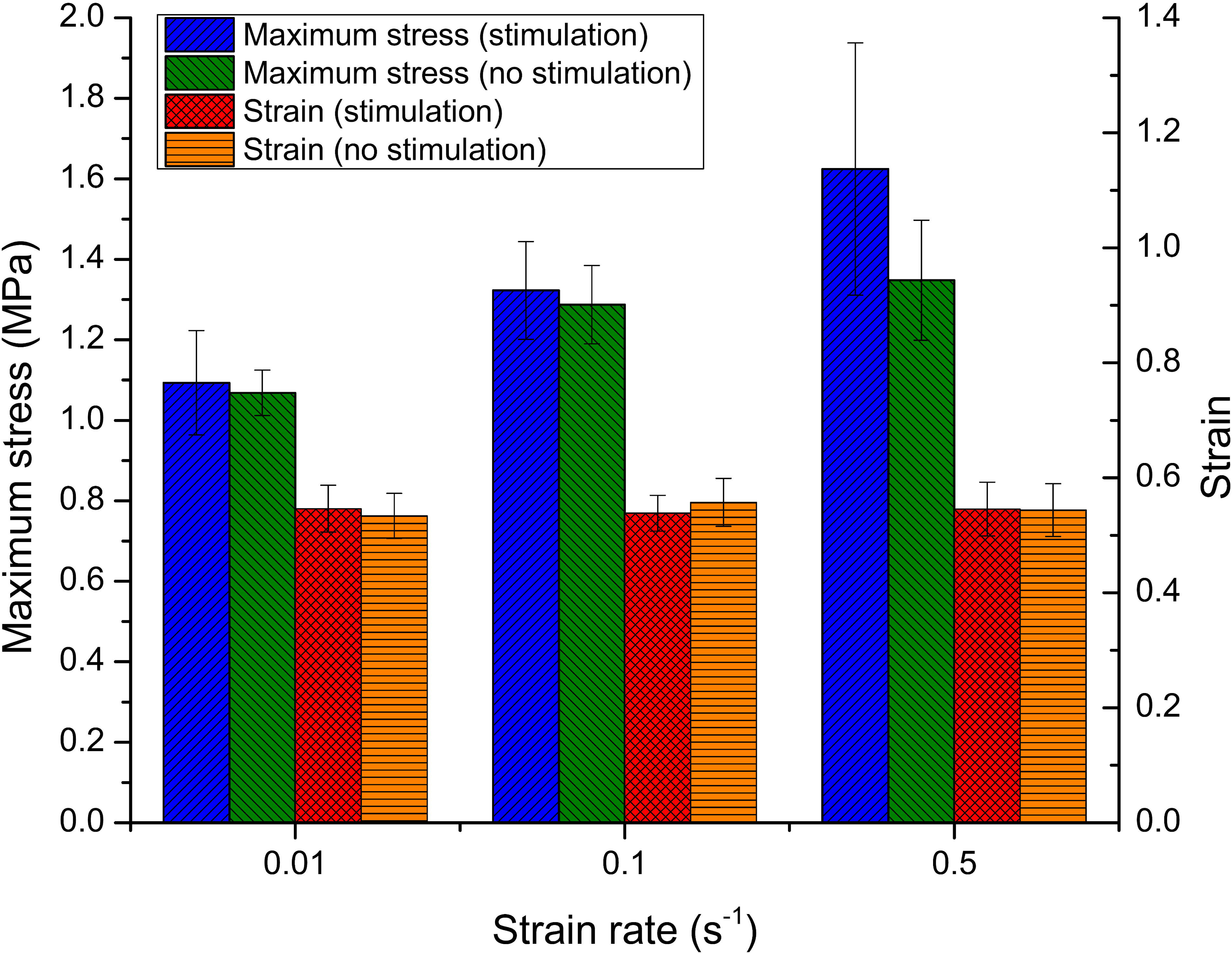
Maximum stress and break strain of samples at low strain rate.

**Fig. 11.**
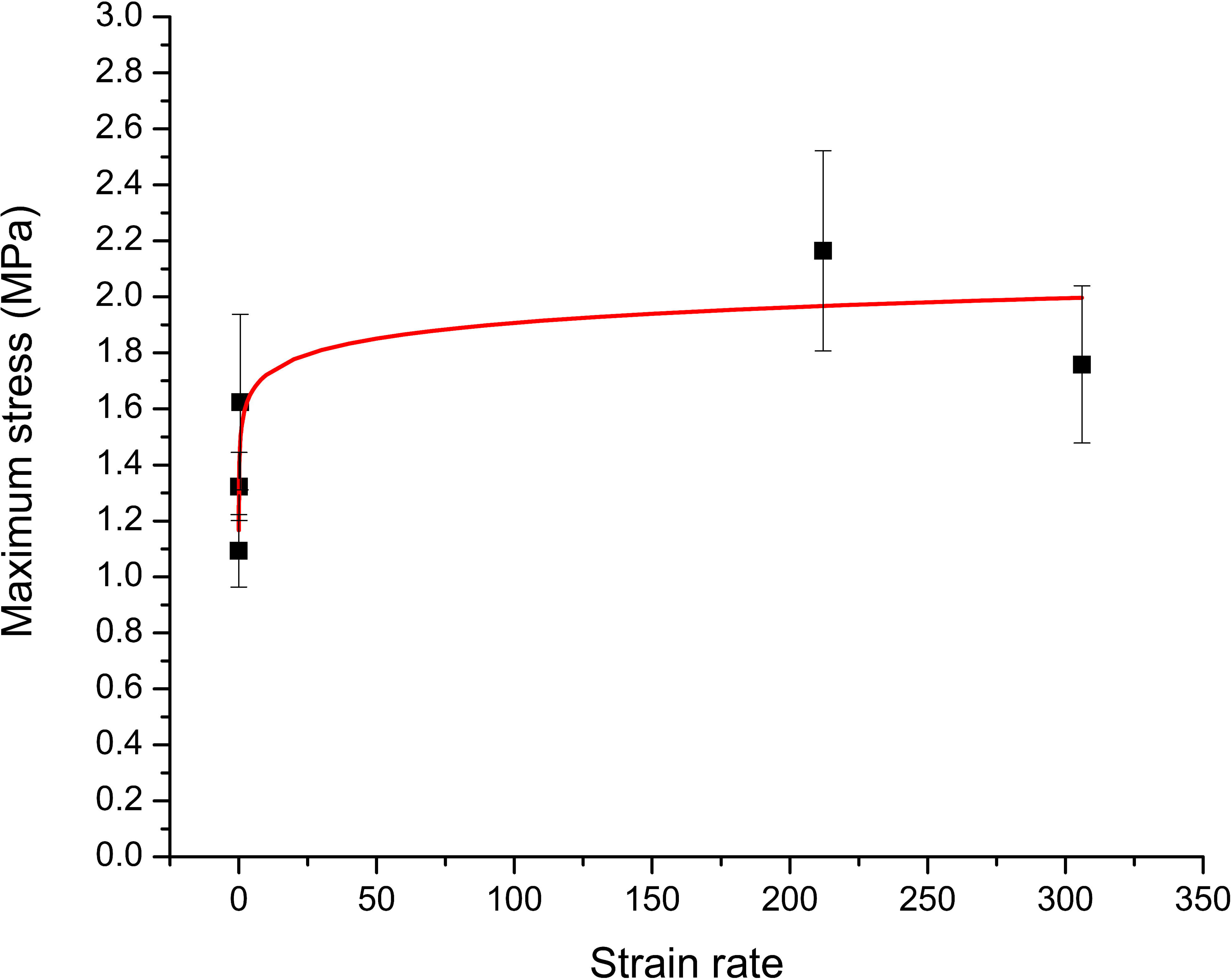
Maximum stress of stimulated samples at various strain rates.

In this research, we also examined the effect of strain and stimulation timing on skeletal muscles’ tensile response at both low and high strain rate ranges. In the low strain rate range, delayed stimulation was applied at 0.2, 0.3 and 0.4 strains during the pulling motion at the strain rate of 0.01 s^-1^. At least nine experiments were performed for each condition, and the moving average curves are shown in Figure 12. When the stimulation was generated at the strain of 0.2, the muscle bundle also stiffened at the beginning, and the maximum stress was similar to samples stimulated at a smaller strain. The stress-strain curve was a sum of active and passive contractions in eccentric contraction. However, samples with stimulations applied at 0.3 and 0.4 strains did not respond similarly. At the strain range of less than 0.3, the stress-strain curve closely followed the unstimulated property curve of the tendon muscle. Nevertheless, the two curves started to separate when the stimulations were applied. Instead of stiffening up, the muscle bundle softened, and the maximum stress decreased drastically. However, maximum stress is reduced by approximately 25% (Figure 13) as the muscle is excited at a higher strain (0.3, 0.4 strain). Conversely, the destructive strain did not change at all experimental conditions (Figures 12 and 13).

**Fig. 12.**
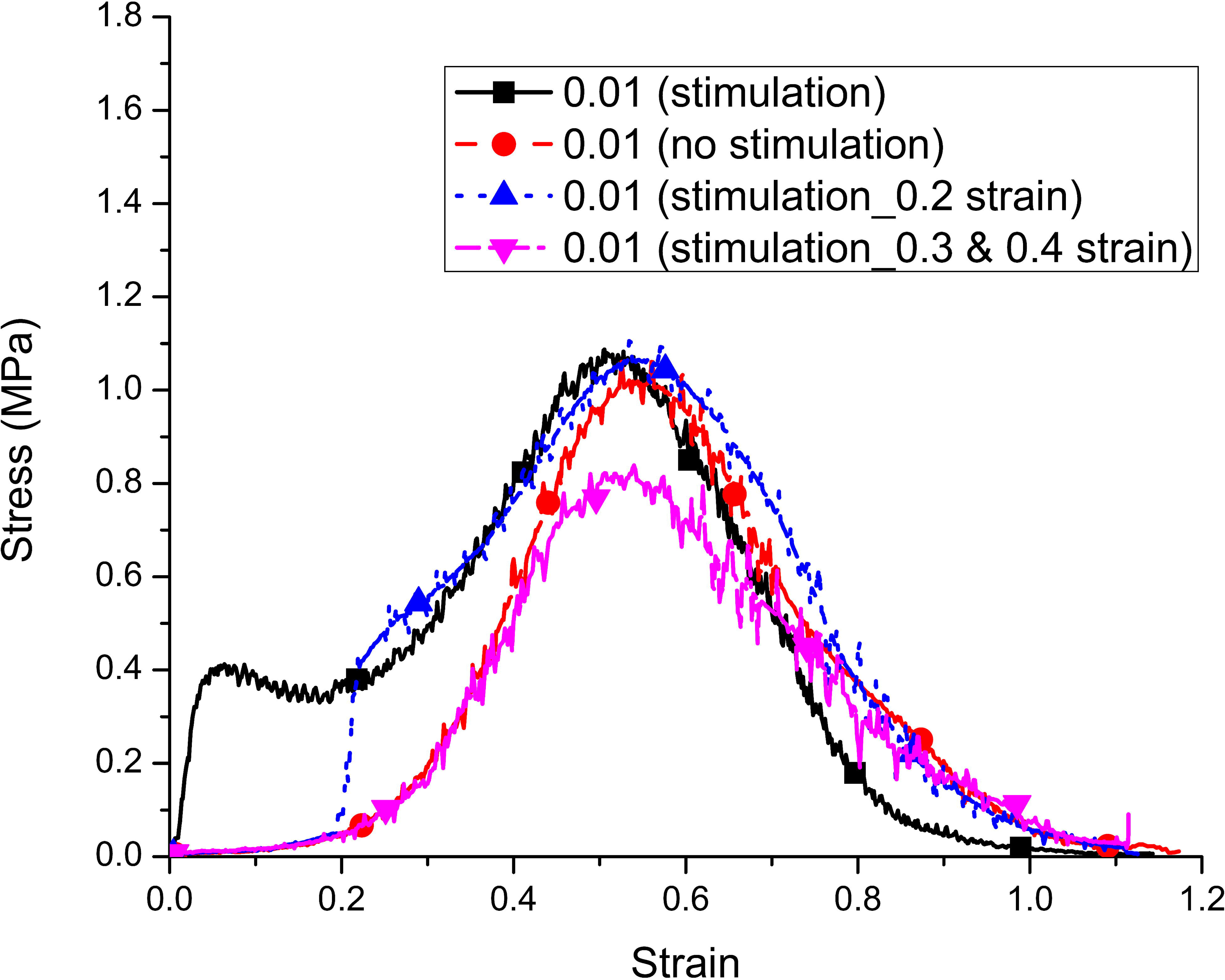
The average stress-strain curves of skeletal muscle at 0.01 strain rate.

**Fig. 13.**
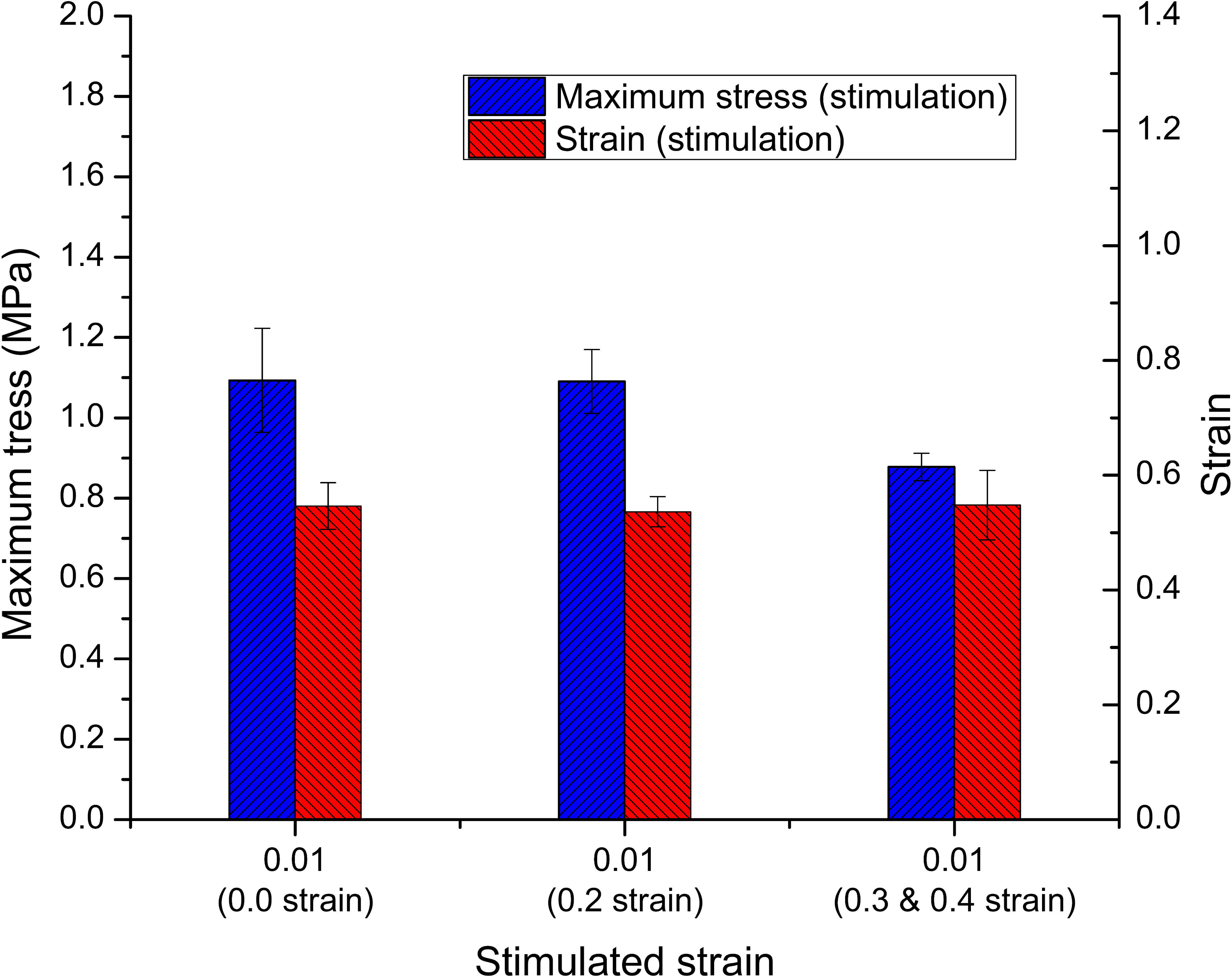
Effect of stimulation strain on maximum stress and corresponding strain at strain rate 0.01 s^-1^.

The role of strain and excitation timing at the high strain rate range was shown in the results of Figures 14 and 15. In Figure 14, the samples were pulled to 10% strain. The sample was then stimulated to produce an isometric contraction in about 3 seconds before being pulled to generate an eccentric contraction. The strain of specimens in Figure 14 was measured in eccentric contraction, and it did not include the 10% original strain created by supporters. Under those experimental conditions, the samples reacted as a linear material at all strain rates from 50 to 300 s^-1^. As the strain rate increased, the elastic modulus still demonstrated a similar decrease as the L0 group, and those values were still higher than Young’s modulus of unstimulated muscle-tendon bundles (Table 2). However, the maximum stress of samples rose and reached about 2.0 Mpa at the strain rate of 150 s^-1^, but tears did not appear. In contrast, the muscle bundles were wholly broken at the tendon-muscle connection area in the strain rates range from 200 s^-1^ to 300 s^-1^, corresponding to a strain from 3.5% to 5% and a maximum stress of 2.0 Mpa. That was similar to the specimens in the first group.

**Fig. 14.**
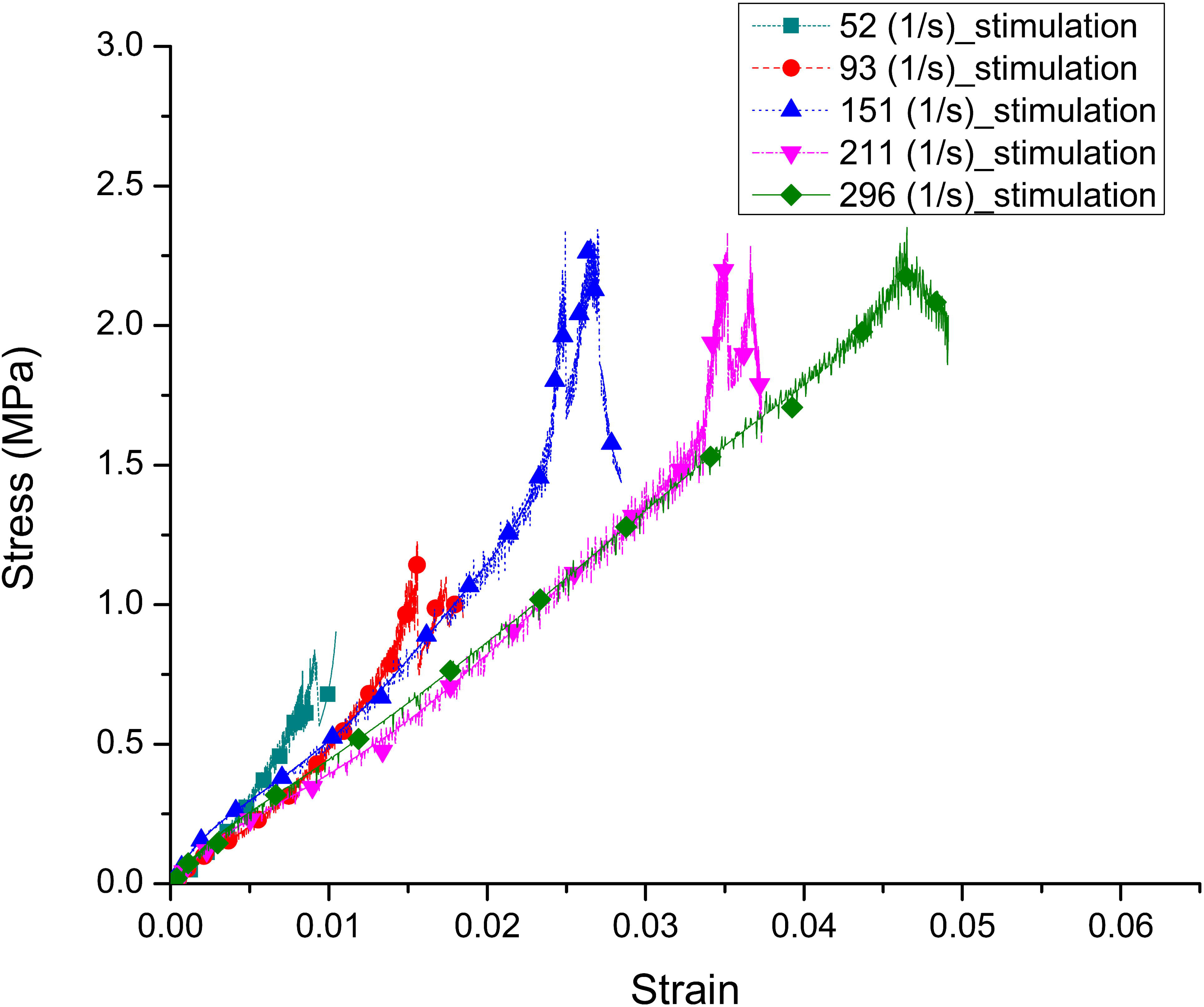
the average stress-strain curves of muscle-tendon specimens stimulated at the 10% length (L10).

**Fig. 15.**
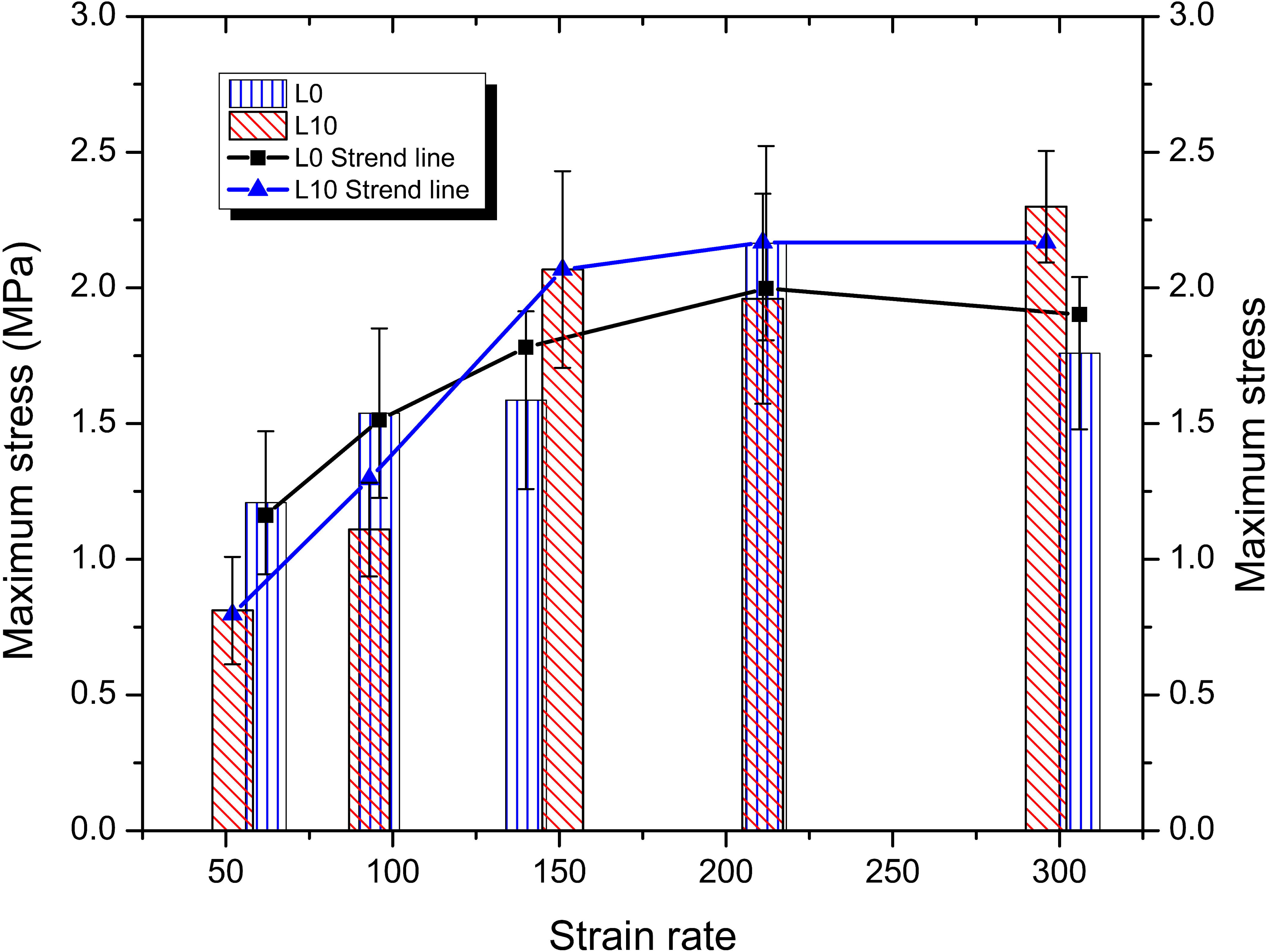
the maximum stress at the various strain rates.

The results in Figure 15 demonstrated that the maximum stress of L0 samples was more significant than L10 samples at strain rates below 150 s^-1^. That could be because the active contraction stress at length L0 was greater than that of the L10 samples. However, the maximum stress in the trend line of the L10 samples was more considerable at the subsequent higher strain rates. This observation showed that the effect of passive stress increased at higher strain and strain rates. That difference was about 0.2 to 0.5 MPa. Although there was a difference in values at various strain rates, both groups followed the same trend: the stress increased as the strain rate increased and remained stable when the strain rate was higher than 150 s^-1^.

## 4. Discussion

The skeletal muscle structure is complex, but sarcomeres linked together by Z-disks are the main building blocks of skeletal muscles [41]. The mechanical properties of muscle bundles at low strain rates are often illustrated by interactions between the sarcomere components, including myosin, actin, and titin [43, 44]. When muscles are applied to an electrical impulse, the sliding between myosin and actin induces muscle contraction, and the resulting stress is called active stress[50–54]. In contrast, the resulting stress is passive under unexcited and stretched muscles. And titin is the main component that determines the passive response of the muscle bundle when under external forces at a low strain rate [41, 55–60]. The response of the muscle-tendon bundle in an eccentric contraction can also be analyzed through the interaction between myosin, actin, titin, extracellular structures, and those two stress components.

The results demonstrated that the muscle-tendon bundle was sensitive to strain rate at both factors, maximum stress and Young’s modulus. Muscle fatigue was not seen from strain rates greater than 0.5 s^-1^. The muscle bundles show a linear property. Young’s modulus increased at low strain rates and decreased in the high strain rate range. While maximum eccentric stress went up and remained at a peak as the strain rate reached 150 s^-1^. On the other hand, the timing of stimulation was also a factor that influenced muscle properties. The maximum stress decreased as the muscles were stimulated at a strain greater than 0.3 (Figure 10). It indicated that a part of the active contractile force had disappeared. That suggested that some of the bonds between myosin and actin were broken, called weak-binding states, and shown by Bernhard Brenner and his colleagues [61–64]. Therefore, in the case of unstimulated and stimulated muscles, a part of the bonds between actin and myosin forms the load stress. We made three assumptions and a Hill-type modelling of the mechanical muscle behavior Figure 16:

1. There were some weak bonds between actin and myosin (which do not generate the contraction) even when the muscles were not stimulated, and those bonds are involved in forming the greatest stress.
2. When the muscle was stimulated, the number of bonds between actin and myosin increased, resulting in active muscle contraction (Figure 8). But this number decreased and maintained the same weak-binding number as unstimulated muscles at high strain where the muscles were destroyed.
3. When the muscles were applied an initiated stimulation at a strain greater than 0.3, new bonds between myosin and actin were not formed (the force of contraction did not increase suddenly), while the original weak-binding bonds were broken by stimulation and large strain [61–64]. That causes the maximum stress to decrease. Original bonds could create about 20 % maximum eccentric stress (Figure 10). These hypotheses supported that the eccentric contraction property of the muscle was determined by the properties of titin, the actin-myosin bond, and the weak-binding state between actin and myosin. This finding demonstrated that load capacity reduces if nerve stimulation is applied late during muscle contractions (at large strain). That may increase the risk of injury. On the other hand, a slow reaction in sports activities perhaps creates a greater risk of injury because the weak bonds are broken.

**Fig. 16.**
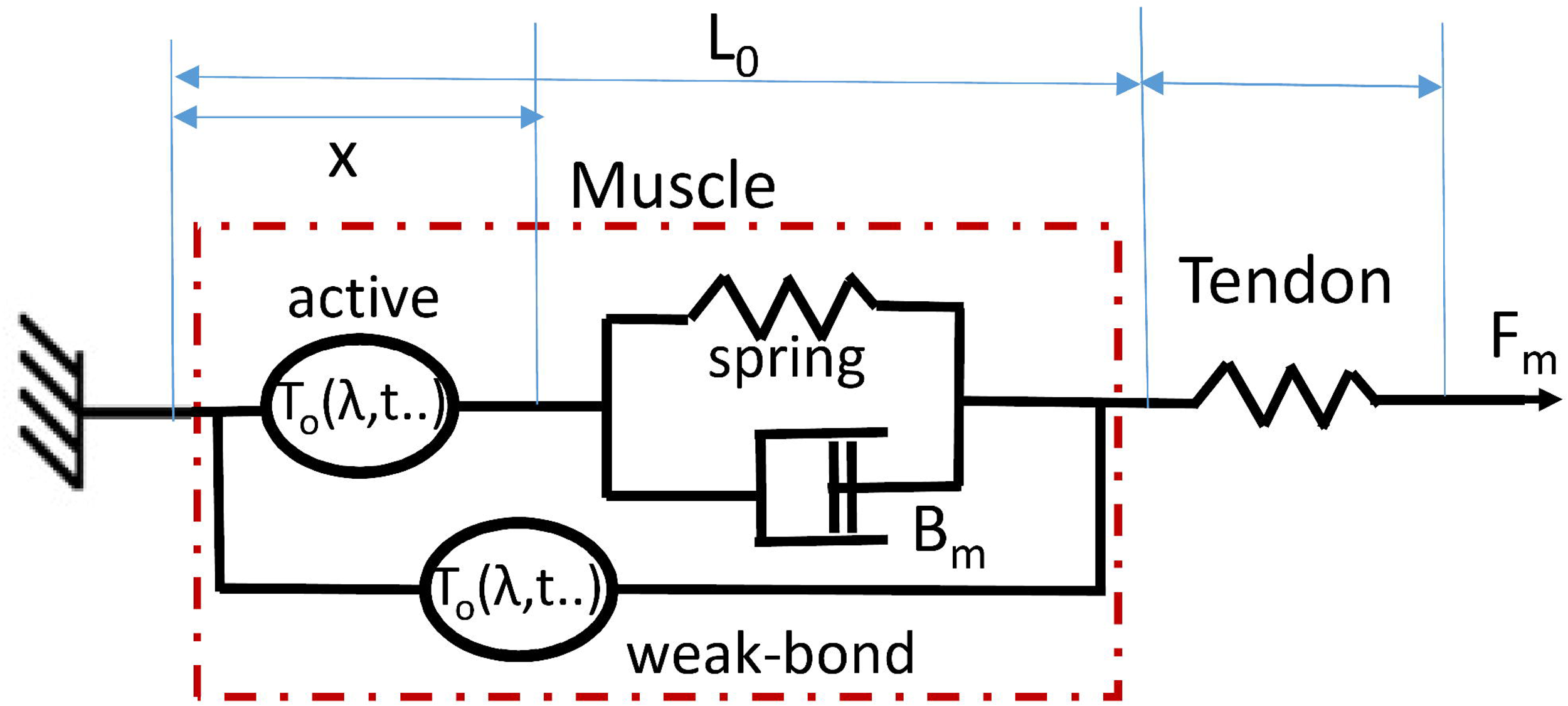
Hill-type modelling of the mechanical muscle behavior.

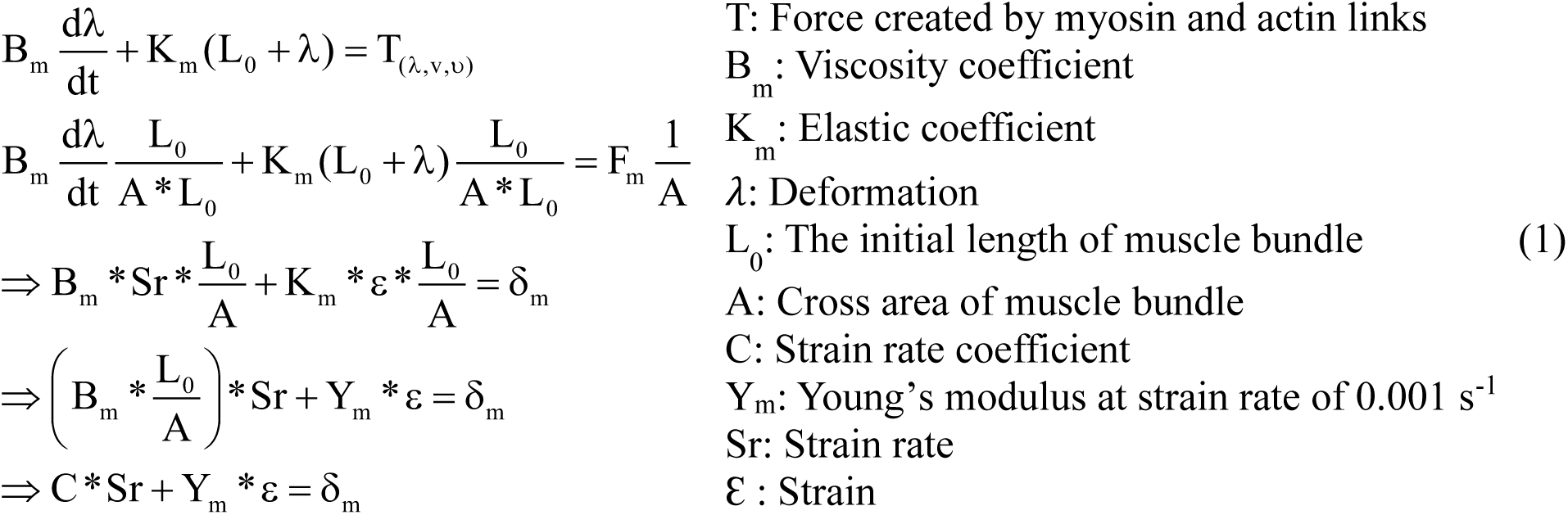

These hypotheses also partly explained the maximum eccentric stress increases phenomenon at the strain rate 0.5 s^-1^ (Figure 7). The rise of eccentric force depending on the contraction velocity was also demonstrated by some research [65, 66]. Because the standard deviation was large, muscle bundles’ maximum stress sometimes dropped to the same breaking force of the unstimulated muscle. That was predicted because the increased breaking force at the stimulated muscle bundles was generated by the unstable bonds between actin and myosin instead of the weak bond [61–64]. These bonds were highly dependent on the properties of the individual muscle bundles. Some studies have also shown that eccentric muscle contraction velocity increased muscle force, but it did not happen at all speeds. That had been suggested that two factors determined it. Firstly, it was because of the superimposed concentric stress [20, 32, 33, 67], which was similar to the effect of excitation frequency occurring at a suitable frequency range. Secondly, it was because muscle fatigue would decrease at a higher speed. However, if the cause were muscle fatigue, the greatest stress increase would occur even at the strain rate of 0.01 s^-1^, but that did not happen. Therefore, the increase in maximum stress has been attributed to superimposed concentric stress. That also illustrates that the risk of injury at high strain rates reduces if the muscles are stimulated. Because the number of actin-myosin bonds is more at the high strain rate, these absorb external force and decrease injury.

Observing and comparing the property results of samples L0 and L10 at high strain rates, they show that the eccentric contraction was sensitive to strain rate. The maximum stress in the L0 samples was more significant than the L10 samples at the two strain rates of 50 and 100 s^-1^. It was predicted because of the effect of active contraction. In the L10 samples, the original length of the samples was larger, so the active muscle contraction force was smaller [26, 68, 69]. That decreased Young’s modulus of the L10 samples; therefore, the maximum stress of those samples went down. In contrast, the maximum stress in sample L10 was greater than sample L0 at the strain rate 100 s^-1^ to 300 s^-1^. Although, the total strain (including deformation before stimulation) of the L10 samples was higher than the L0 samples, and the active stress generated in the L10 samples was smaller than in the L0 samples. It showed that the role of active contractile force has decreased, and the role of passive contractile force has increased at 130 s^-1^ high strain rates [65, 66, 70]. In other words, the bonds between myosin and actin that conferred active stress could reduce in number at high strain rates, and the titin and extracellular structures played a more significant role at high strain rates[71–73]. That could explain why Young’s modulus of stimulated muscle-tendon bundles was higher than the unstimulated samples, but Young’s modulus decreased as the strain rate increased at the high strain rate ranges in L0 and L10 groups. As the role of the links between myosin and action was significantly reduced at the strain rate of 300 s^-1^, Young’s modulus of the excited samples was reduced to close to that of the unexcited samples.

In addition, the maximum stress value in both groups of samples L0 and L10 at strain rates of 100 and 150 s^-1^ was higher than 20% of failure stress in the stimulated samples tested at the 0.5 s^-1^ strain rate. However, the samples did not show fractures. That has shown that stress was not the only parameter affecting muscle injury. It has been confirmed that strain rate is the more influential factor [11, 22]. Conversely, the maximum stresses in both samples at the strain rate of 200 s^-1^ did not change significantly, and the strain only increased by 1% compared to the 150 s^-1^ strain rate (from 2.5 to 3.5%). However, the samples were completely broken. That raised a question: the change in strain rate or the small change in strain caused the muscle rupture in this case. We hypothesized that the muscle region did not have enough time to stretch and absorb energy when pulling at high strain rates. At the same time, the instantaneous load capacity of the muscle area was large at a high strain rate [65, 66]. So the deformation of the muscle bundle was focused on the tendon area. That caused the rupture in the tendon muscle region, and previous studies have shown it has the highest injury rate in the muscle-tendon bundle [49, 74]. While the injury position was a musculotendinous junction as the samples were stimulated and pulled at a low strain rate (our paper).

Another essential factor to be considered here was that the strain of L0 tested at strain rates of 200 and 300 s^-1^ was about 3.5%, but the samples were broken. In addition, the L10 samples were stretched by 10% (the maximum strain of the hamstring muscle during sprinting [75]) before an eccentric contraction was applied. Therefore, the total strain (including deformation before stimulation) of L10 samples at strain rates below 150 s^-1^ was still higher than the L0 samples at strain rates above 200 s^-1^_;_ however, L0 samples were completely torn. Nevertheless, the L10 samples at strain rates below 150 s^-1^ were not broken, and the samples only showed fractures at strain rates of 200 s^-1^ and 300 s^-1^, similar to the L0 samples. Therefore, it was proposed that a 10% increase in sample strain before the samples had an eccentric contraction at a high strain rate did not significantly affect injury. That could also be explained by the hypothesis above. Where deformation of the tendon-muscle bundle had occurred in tension at the high strain rate would focus on the tendon area to be more deformed and cause injuries. Therefore, studies suggesting that deformity affects muscle injury more should consider this factor.

This study also showed a difference between stimulated and unstimulated tendon muscle bundles. Only a few unstimulated samples suffered minor damage, while all stimulated samples were completely broken at a strain rate higher than 200 s^-1^. That showed that electrical impulse stimulation increased the risk of muscle-tendon bundle injury at high strain rates because the stimulus increased Young’s modulus of the muscle bundles and increased the deformation of the tendon area, thereby increasing the risk of injury at a high strain rate.

Our study aimed to evaluate the response and injury risk of the muscle-tendon bundle at various strain rates and stimulating times. In addition, the deformation of the muscle bundle is also a factor that we were particularly interested in this study. The results indicate that the eccentric-contraction property of the tendon-muscle bundle is sensitive to strain rate. The maximum stress increases with the rise of strain rate and remains at the peak of about 2 MPa. Young’s modulus illustrates a rise in the lower strain rate ranges than 200 s^-1^, but it reduces at the higher strain rates. Injuries often occur at the musculotendinous junction due to muscle strain. The injuries concentrate in the tendon area with very small deformation, only about 3.5 to 5% when the strain rate is higher than 150 s^-1^. That is because the load capacity of the muscle is higher while the absorbed energy is reduced, which causes the increase of deformation in the tendon area. In addition, stimulation also influences the risk of injury at high strain rates because it increases stiffness in the muscle area, which causes a higher strain in the tendon region. While deformation of about 10%, before muscle-tendon bundles are pulled at high strain rates, did not significantly affect the risk of injury.

The timing of muscle stimulation, which is determined through muscle strain, also causes a change in the load capacity of the tendon bundle. When the stimulus is generated at a strain of less than 0.3, the maximum stress of the muscle bundle does not change. Nevertheless, when stimulations are produced at a strain greater than 0.3, a reduction of about 20% of the maximum stress is observed. Therefore, it is predicted that some bonds between myosin and actin persist even when the muscle bundles are not stimulated. These links will participate in the deformation process of the muscle and create the largest part of the stress. However, when the muscle bundles are stimulated at considerable strain, these connections will be broken and cause the most significant stress reduction [61–64]. That showed that the time of stimulation also influenced muscle injury.

## Acknowledgments

This work was supported by the Ministry of Science and Technology, Taiwan, MOST 109-2637-E-992-011.

## CRediT authorship contribution statement

Dat Trong Tran: Conceptualization, Methodology, Investigation, Writing - original draft, Review & editing.

Liren Tsai: Methodology, Review & editing, Supervision, Validation, Corresponding author.

